# Signal space beamforming for gain- and geometry-independent interference suppression in wearable MEG

**DOI:** 10.64898/2026.01.22.700798

**Authors:** Maxime Ferez, Pierre Corvilain, Odile Feys, Chiara Capparini, Laureline Fourdin, Julie Bertels, Xavier De Tiège, Vincent Wens

## Abstract

Optically pumped magnetometers (OPMs) for wearable magnetoencephalography (MEG) offer substantial flexibility compared to cryogenic MEG, but they also introduce new challenges related to environmental noise and movement artefacts. State-of-the-art denoising techniques such as signal space separation require not only dense OPM arrays but also the fine calibration of sensor gain and geometry. This may complicate their use in complex experimental situations involving uncooperative patients or subjects such as newborns or fetuses, where OPM positioning may be unstable or plain unknown. To address this limitation, we introduce signal space beamforming (SSB), a version of beamforming applied to sensor signals and designed to suppress interferences independently of sensor gain and geometry. We show that SSB operates via a trade-off between noise suppression and neural signal preservation that is controlled by a single soft-threshold parameter, which we calibrated using phantom measurements. We then validated the effectiveness of SSB across several datasets. Using interictal OPM-MEG recordings in epileptic patients, SSB successfully cleaned signals while preserving epileptiform discharges. Moving onto cutting-edge early neurodevelopmental OPM-MEG recordings, SSB allowed to recover auditory evoked responses in newborns and fetuses similar to previous findings but with more streamlined preprocessing and higher response amplitude in the case of the fetal data. We contend that SSB provides a pragmatic solution to exploit OPM-MEG data recorded in challenging conditions where geometry-dependent methods cannot be used, opening new frontiers for pioneering clinical and fundamental applications of the OPM technology.

## 1. Introduction

The development of optically pumped magnetometers (OPMs) represents a technological advance that could arguably transform the field of magnetoencephalography (MEG) (Boto et al., 2018; Holmes et al., 2023b). The main practical advantage of OPM-MEG is the flexibility with which OPMs can be arranged, as opposed to the rigidity of conventional cryogenic MEG helmets. This has initiated sophisticated neurocognitive and clinical applications that go beyond what cryogenic MEG allows. Wearable systems have enabled neuromagnetic activity recordings in natural settings (Rivero et al., 2025; Seymour et al., 2021)—including ambulatory movements (Holmes et al., 2023c) and fully portable miniaturized systems (Schofield et al., 2024)—as well as recordings of spinal and peripheral activity (Bu et al., 2022; Mardell et al., 2024). They have allowed the recording of epileptiform discharges both between (Feys et al., 2022, 2023e, 2023c, 2025b; Hillebrand et al., 2023; Schwartz et al., 2025) and during (Feys et al., 2023d) seizures in infants (Feys et al., 2023a), children (Feys et al., 2022), and adults (Pedersen et al., 2022) with epilepsy. Further, OPM-MEG has paved the way for sensory and neurocognitive investigations covering the lifespan (Hill et al., 2019) down to infants (Corvilain et al., 2025b) and even prenatal life (Corvilain et al., 2025a).

In wearable OPM-MEG settings, head movements—and more generally variations in the position and orientation of individual sensors moving in the environmental magnetic field—translate into signal artefacts that substantially interfere with OPM recordings and lower their signal-to-noise ratio (SNR). Interference suppression is therefore a major theme of OPM-MEG methodology (Seymour et al., 2022). One fundamental aspect of OPM-MEG signal denoising is the hardware nulling of environmental fields using a combination of passive and active magnetic shielding (Boto et al., 2018; Holmes et al., 2019). Another is the implementation of software denoising techniques, two of which have proven exceedingly impactful in cryogenic MEG and have been adapted to OPM-MEG. First, signal space separation (SSS) suppresses environmental magnetic interferences directly at the sensor level on the basis of truncated multipolar expansions (Taulu et al., 2004). Variations specific to OPM-MEG have been developed such as the homogenous field correction (Tierney et al., 2021), the iterative SSS (Holmes et al., 2023a), adaptive multipole modeling (Tierney et al., 2024), and the multi-origin SSS (McPherson et al., 2025). Second, beamforming has been largely used as a source projection technique that intrinsically denoises MEG signals at the source level, by suppressing correlated activity typical of external noise (Hillebrand & Barnes, 2005; Van Veen et al., 1997). In the absence of principled sensor-level denoising techniques for OPM signals, beamforming has been argued to provide efficient interference suppression (Boto et al., 2021; Seymour et al., 2022). Nevertheless, these two techniques require a fine knowledge of sensor gain (i.e., the factor converting voltage recordings into magnetic signals) and geometry (i.e., relative positions and orientations)—a.k.a. MEG calibration. The accuracy of this calibration controls that of the multipolar expansion underlying SSS (Nurminen et al., 2008) and of the MEG forward model underlying beamformer source projection (Lucena Gómez et al., 2021; Steinsträter et al., 2010). The flexibility of OPM-MEG poses a challenge to apply these denoising methods, since OPM gain and geometry must be calibrated on a per-recording basis and could even change during a recording, in contrast with cryogenic MEG where this measurement can be done once in factory. Hardware solutions range from optical scanning (Zetter et al., 2018, 2019) to the recent development of bespoke coil systems (Hill et al., 2025; Iivanainen et al., 2022). Still, there are situations where OPM gain and geometry are difficult to calibrate accurately in practice. This applies for example to new OPM-MEG laboratories that do not have yet access to these techniques, but also—and perhaps more critically—to recordings performed in participants with poor cooperation. Examples encompass patients with severe forms of epilepsy, neurodevelopmental or degenerative disorders, as well as newborns and even fetuses in the field of early neurodevelopmental MEG. These cases, on top of being intrinsically challenging, are plagued by the difficulty to apply powerful denoising techniques.

We introduce here a novel interference suppression strategy that leverages the intrinsic denoising property of beamformers, but with an adaptation that allows to work at the sensor level independently of OPM calibration. The technique—coined signal space beamforming (SSB)— eliminates highly correlated sensor-level activity characteristic of environmental noise and movement artefacts that typically dominate over neural contributions in OPM-MEG signals. Since brain activity also involves correlations across sensors, SSB must operate with a trade-off between noise suppression and preservation of the neural activity of interest. We demonstrate that SSB depends on a single free parameter that controls the threshold between suppression and preservation. We calibrated the SSB parameter experimentally using empty-room OPM measurements on a phantom head and subsequently probed several aspects of SSB accuracy with a series of OPM experiments. Sheer denoising quality was assessed using noise recordings under controlled external disturbances that reproduced different types of OPM artefacts. Preservation of neural activity after SSB denoising was assessed using recordings in epileptic patients (Feys et al., 2022, 2025b). We used here interictal epileptiform discharges (IEDs) as a model of neural activity with a clearly recognizable signature. Finally, we moved onto cutting-edge early neurodevelopmental data for which performing OPM-MEG calibration had not been possible (Corvilain et al., 2025b, 2025a). The application of SSB to fetal OPM-MEG data is of particular interest, since the convergence of SSB results with the previously found responses—obtained after a convoluted signal processing (Corvilain et al., 2025a)—would corroborate the neural origin of fetal OPM-MEG responses.

## 2. Theory

This section outlines the theoretical foundations of interference suppression by SSB. In essence, SSB corresponds to a classical unit-gain linearly constrained minimum variance beamformer (Sekihara & Nagarajan, 2008), but with the adaptations that the MEG forward model is replaced by the identity matrix and that the data covariance matrix is appropriately regularized. This effectively replaces channels by “virtual channels” with a preserved gain, but in which correlated activity of large amplitude is suppressed. The amplitude threshold, above which correlated activity is suppressed and below which it is preserved, is controlled by the covariance regularization parameter. We briefly develop here the mathematics of SSB. The main practical outcome is the SSB spatial filter equation implementing SSB interference suppression. We also highlight the key role of the amplitude threshold, the main assumptions behind OPM-MEG interference suppression with SSB, and a heuristic strategy to calibrate the threshold parameter.

### 2.1 The SSB spatial filter equation

We consider a MEG system composed of a number *N* of channels recording magnetic signals *b*_*i*_(*t*) over time *t* (with channels indexed by *i* = 1, …, *N*), which we shall gather into *N* × 1 vector signals ***b***(*t*). The SSB corresponds to a spatial filter 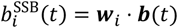 with 1 × *N* weights ***w***_*i*_ defined by the minimization of output data variance Var 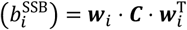 under the unit-gain constraint (***w***_*i*_)_*i*_ = 1. Here ***C*** = cov(***b***) denotes the *N* × *N* input data covariance matrix. This classical beamformer problem leads to the solution (***w***_*i*_)_*j*_ = (***C***^−1^)_*ij*_/(***C***^−1^)_*ii*_ (Van Veen et al., 1997). In general, inversion of the matrix ***C*** may be ill-conditioned and require regularization beforehand. We chose here Tikhonov regularization where ***C*** is replaced by 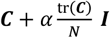, with ***I*** denoting the *N* × *N* identity matrix, tr the trace, and *α* the regularization parameter measured as a percentage of the mean input variance tr(***C***)/*N*. Taken together, these considerations lead to the main SSB equation

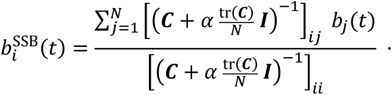

Importantly, the choice of the free parameter *α* in the context of SSB denoising is not merely about regularization of matrix inversion but represents a full-fledged part of interference suppression. We continue with some theoretical developments that justify this claim. Readers willing to focus on the practical implementation of the SSB (including experimental calibration of the parameter *α*) and on its evaluation as an OPM denoising algorithm may proceed directly to the experiment Sections 3 and 4.

### 2.2 The SSB preserves sensor gain

A first key property is that virtual channels 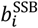 possess the same gain as the input channels *b*_*i*_. One way to measure sensor gain would be to arrange the *i*^th^ channel *b*_*i*_ to sense, e.g., a calibrating sinusoidal magnetic field of 1 nT amplitude, while keeping all other channels insensitive to it (i.e., *b*_*j*_ = 0 for *j* ≠ *i*). In such a situation, there is no cross-channel correlation so the covariance ***C*** is diagonal and the SSB equation leads to 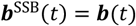, i.e., the SSB spatial filter does not modify sensor response to a calibration field. In other words, sensor gain is preserved.

### 2.3 The SSB suppresses correlated activity with large (but not small) amplitude

The general property of beamformers that correlated activity is suppressed (by means of output data variance minimization; see (Van Veen et al., 1997)) translates for the SSB into the suppression of *any* activity showing cross-channel correlations, whether due to external noise or generated by neural dipolar activations. This would *a priori* be a huge flaw of the approach; however, this suppression effect turns out to depend on the amplitude of the correlated activity.

As a minimal toy model to illustrate this, let us consider the situation of an idealized signal artefact ***a***(*t*) superposed on an independent neural signal ***n***(*t*), so ***b***(*t*) = ***n***(*t*) + ***a***(*t*). Our idealized artefact involves only two channels, *i* and *j*, with a perfect correlation between them and a large variance 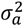. The neural signal is distributed across channels with lower variance 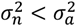. To simplify the analysis as much as possible, we momentarily set *α* = 0, effectively allowing to discard cross-correlations in the neural signal (see next subsection). The corresponding covariance matrix is 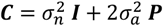, where ***p*** is the *N* × *N* projection matrix onto the artefact subspace (i.e., its only non-zero entries are *p*_*ii*_ = *p*_*jj*_ = *p*_*ij*_ = *p*_*ji*_ = 1/2), and the inverse of ***C*** is

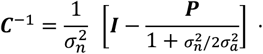

Therefore, the SSB spatial filter outputs a signal 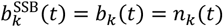 that is left unmodified for non-artefactual channels *k* ≠ *i, j* while modifying the artefactual channel *b*_*i*_(*t*) = *n*_*i*_(*t*) + *a*(*t*) into

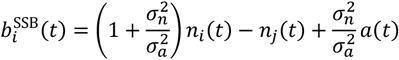

and similarly for channel *j*. This expression reveals that the suppression of the artefact signal *a*(*t*) is modulated by its amplitude σ_*a*_ relative to the amplitude σ_*n*_ of the rest of the signal, in such a way that suppression is more efficient for large-amplitude artefacts. In the situation of a dominating artefact 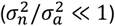, these channels become 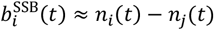 and 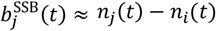, reflecting that virtual sensors are artefact-free and orthogonalized with respect to the artefact subspace. This illustrates the fact that SSB suppresses correlated artefacts only if they are of large amplitude, which is precisely the situation we target in OPM-MEG.

### 2.4 The SSB parameter controls the threshold between suppression of high-amplitude signals and preservation of low-amplitude signals

As last piece of theory, let us reformulate the effect of SSB spatial filtering, this time not in terms of channel signals but in terms of modes of correlated MEG activity as obtained by principal component analysis (PCA). This allows to generalize the above notion that suppression of correlated signal artefacts requires a separation of amplitudes but also characterizes how the parameter *α* relates to that separation.

Specifically, input data are now viewed as a combination of different modes *m* = 1, …, *N* according to the eigen-decomposition 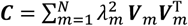 (i.e., principal components). The eigenvalue 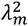 measures the variance of the *m*^th^ mode, the unit *N* × 1 eigenvector ***V***_*m*_ characterizes its topography, and its temporal activity is encoded in the standardized time course 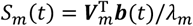. Further noting that tr(***C***)/*N* corresponds to the mean eigenvalue ⟨*λ*^2^⟩, the SSB spatial filter equation becomes

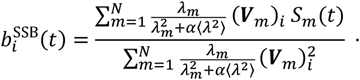

This reformulation connects SSB to a “soft” version of PCA denoising (whereby a fixed number of the highest-variance modes are removed altogether based on a hard threshold on 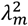, instead of being smoothly weighted down). A critical advantage of SSB is that it preserves sensor gain.

Similarly to the toy model, the above expression describes how SSB suppresses modes of correlated activity with large relative variance in the sense 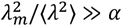, and thus reveals that the parameter *α* plays the role of a (soft) threshold for this suppression. In other words, the SSB threshold parameter *α* determines which correlated activity is well suppressed and which is preserved.

### 2.5 Interference suppression with SSB requires a heuristic calibration of its threshold parameter

Onto the practical context of OPM-MEG denoising, the rationale behind the SSB technique is that typical environmental and movement-related OPM noises correspond to correlated signal artefacts with large amplitudes (i.e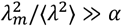, formally defining an “interference subspace”). Neural activity of interest is characterized by substantially smaller amplitudes (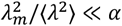, defining a “signal subspace”). In a sense, SSB interference suppression assumes that there is a well-defined gap in amplitude separating the interference and signal subspaces. The threshold parameter *α* should be calibrated to lie within this gap; this is the first step towards a practical implementation of SSB.

Since this gap is *a priori* unknown, we need some heuristic approach. The interference subspace targeted by SSB can be characterized using noise OPM-MEG recordings. Most of their variance lies within the interference subspace, though a subdominant fraction of variance also probes the complementary subspace. This gap structure leads to an ill-conditioned covariance whose inverse will exacerbate the contribution of the small-variance complement to the interference subspace, leading to spuriously high output signal variance after SSB. Our heuristic is to calibrate *α* to the smallest value that minimizes the output variance for OPM noise recordings, so as to suppress the interference subspace while preserving the complement subspace, which—in OPM-MEG recordings containing brain activity—should cover the signal subspace.

## 3. Materials and Methods

### 3.1 System and data acquisition

For all acquisitions described below, OPM-MEG recordings were performed using alkali quantum zero-field magnetometers (Gen2/Gen3 QZFM, QuSpin Inc.) housed in custom 3D-printed holders sewn on flexible caps for phantom head, epilepsy, and neonate recordings (Feys et al., 2022, 2025b; Hill et al., 2020) or belts for fetal recordings (Corvilain et al., 2025a). The number of OPMs used varied from experiment to experiment (see below). All experiments took place in a magnetic shielded room (MSR; OPM-optimized Compact MuRoom, Magnetic Shields Ltd.). The magnetic field within the room was reduced by degaussing the MSR (Altarev et al., 2015) and by energizing wall coils (MuCoils, Cerca Magnetics; coil currents based on a mapping of static background field), leading to a residual magnetic field below 1–2 nT within a 40-cm-wide cubic region of interest. Analog OPM signals were acquired using a default gain of 2.7 V/nT and digitized at 1200 Hz sampling frequency via a 16-bit digital acquisition device (National Instruments) operated with a custom Python-based acquisition control program.

### 3.2 Data preparation and denoising

Data preprocessing and analysis were performed with custom-made scripts for MATLAB (Version R2022b, MathWorks Inc.) Raw OPM data were first inspected visually for the identification and exclusion of flat or excessively noisy channels, for the identification of noise peak frequencies arising in the power spectrum that were removed by notch filters, and for the marking of bad time segments. An independent component analysis (ICA; (Vigario et al., 2000)) was also applied to isolate and remove heartbeat artefacts (FastICA algorithm with *tanh* nonlinearity (Hyvarinen, 1999)).

Interference suppression based on SSB was then directly applied to these data according to our main spatial filter equation (Theory section 2.1). The covariance matrix ***C*** entering the SSB spatial filter was computed without further frequency filtering of input data (but excluding bad time segments). The SSB threshold parameter α was calibrated using empty-room phantom recordings described below. To assess the impact of SSB on subsequent analyses, we compared results with the same data without any further denoising, and the same data denoised using dedicated pipelines based on PCA (with 3 or 15 principal components removed, as previously applied in the data considered here; see (Corvilain et al., 2025b; Feys et al., 2022, 2025b)) and regression of external signals (Corvilain et al., 2025a).

### 3.3. Calibration of the SSB threshold parameter

Calibration of the threshold parameter α (Theory section 2.1) was based on a practical implementation of the heuristic outlined in Theory section 2.5. We recorded 5 minutes of emptyroom signals with 53 OPMs (30 Gen2 biaxial OPMs, 23 Gen3 triaxial OPMs) on a phantom head with a flexible cap. The phantom was located within the 40-cm-wide recording cube associated to the sitting position. After removing flat and noisy channels, 47 OPMs (26 Gen2, 21 Gen3) were kept for analysis. Data were preprocessed and denoised using SSB, increasing prospective values of α ranging from 0% to 5% with 1% increments and from 5% to 50% with 5% increments. In each case, denoising efficiency was assessed by considering signal power in the 1–40 Hz frequency range (to which the epilepsy and neurodevelopmental data analyses below were limited), and by comparison to the same data without SSB denoising. Signal power was obtained from the power spectral density (PSD, computed using 10-second-long windows free from bad segments and averaging across all windows) averaged across channels and across frequencies between 1 and 40 Hz. The calibration heuristic consisted of selecting the smallest of the prospective values α beyond which signal power did not decrease further. The resulting calibrated SSB threshold was used in all the subsequent experiments described in the next sections.

To assess reproducibility of this calibration over time, we repeated the same procedure with empty-room signals recorded 9 months later, twice a day for 5 consecutive days. We used 45 OPMs on the phantom head (11 Gen2 biaxial, 10 Gen3 biaxial, 24 Gen3 triaxial), of which 42.4 ±

2.3 (mean ± SD over 10 recordings) were retained after flat/noisy channel exclusion (9.5 ± 2.1 Gen2 biaxial, 10 Gen3 biaxial, 22.9 ± 0.7 Gen3 triaxial). We used yet another phantom head in the lower position corresponding to fetal recordings (with potentially different remnant environmental magnetic field) to check the validity of this calibration for fetal OPM-MEG data.

### 3.4 Further assessing SSB denoising performance

The above-mentioned SSB calibration step makes the implicit assumption that environmental magnetic field fluctuations captured by the empty-room OPM recording faithfully sample the interference subspace (Theory section 2.5). The rationale is that other artefacts (such as head movements, sensor vibrations, etc.) relate to OPMs moving in the very same field; their temporal dynamics are different, but it is plausible that spatial patterns of cross-channel correlations are similar. As proof-of-concept of this idea, we considered several recordings, both simulating different types of OPM noises on a phantom head and using physiological noise from a human subject. In all cases, we quantified the denoising performance of calibrated SSB as above, using the mean 1–40 Hz PSD comparatively to data without interference suppression.

#### Large field perturbations

We recorded OPM signals on the phantom head for 35 seconds, while pulsing successively the 22 wall coils of the MSR in turn. This generated strong but relatively homogeneous fields inside the MSR. Of note, during this test the coils were not available for active compensation of the environmental field in the MSR, which was thus larger in this experiment. The flexible cap was prepared with 53 OPMs, 46 of which were kept for analysis after flat/noisy channel exclusion (25 Gen2 biaxial, 21 Gen3 triaxial). ***Head movement artefacts***. We simulated head movements (i.e., rigid translations and rotations common to all sensors) by recording the empty-room OPM signals while applying a gentle movement to the phantom head, for a total duration of 1 min. We used 43 OPMs (23 Gen2 biaxial, 20 Gen3 triaxial); 33 were retained for analysis (16 Gen2, 17 Gen3). ***Vibration artefacts***. We induced OPM vibrations (i.e., non-rigid sensor movements independent across sensors) by submitting loud sounds (85 dB intensity at the helmet) delivered to the phantom head using a MEG-compatible flat speaker (SSH sound shower, Panphonics). The sound sequence lasted 5.25 min and consisted of 500 Hz pure tones (100 ms duration including 15 ms of fade-in and fade-out, with inter-stimulus interval of 600 ms ± 100 ms) generated using a custom Python code. Such non-rigid OPM movement artefacts are of interest since they occur with flexible caps during head movements (contrary to rigid helmets; see (Hill et al., 2020)). The goal of this experiment was to assess the ability of SSB to deal with such artefacts. These sound vibration artefacts were dealt with in a previous OPM-MEG study (Corvilain et al., 2025a) using a linear regression technique at the level of auditory evoked response fields (ERFs). We thus supplemented our PSD analysis with a direct ERF analysis of sensor vibrations locked onto the time of sound presentation (band-filter, 1–40 Hz; epoch, 100 ms pre-stimulus to 700 ms post-stimulus, baseline correction over 100 ms pre-stimulus, averaging over all epochs without any bad segment). We used 53 OPMs, 6 of which were rejected (26 Gen2 biaxial, 21 Gen3 triaxial). ***Physiological artefacts***. We then recorded typical artefacts encountered during acquisition with a subject. We measured a healthy participant (35 years, male) wearing a flexible cap with 51 OPMs (29 Gen2 biaxial, 23 Gen3 triaxial; 46 kept for analysis, 26 Gen2, 20 Gen3) while he performed a sequence of specific actions, each held for approximately 20 seconds. The protocol included, in order: eye blinking, lateral and then horizontal eye movements, contraction of jaw muscles, clenching of the teeth, contraction of the neck muscles, and lastly free head movements.

### 3.5 Assessing the preservation of neural activity after SSB

While the above experiments examined the ability of calibrated SSB to suppress typical OPM interferences, they did not allow to assess whether neural activity is preserved. Neural activity in the signal subspace exhibits cross-channel correlations so there is no guarantee that SSB does not suppress it too, either because of failure of our heuristic calibration procedure or because of the invalidity of the hypothesis of a gap separating the interference and the signal subspaces (Theory section 2.5). To investigate this question, we analyzed resting-state data from epileptic patients with spontaneous IEDs, which we use here as a model of neural activity with a high SNR. Demonstration that large-amplitude neural signals are preserved after SSB implies preservation of lower amplitude neural signals too, because the SSB threshold α lies on a linear axis of signal variances (Theory section 2.5).

We selected 6 patients with large and frequent IEDs from previous OPM-MEG studies. Four patients were children (age: 9 years ± 2.8 years, mean ± SD over patients) suffering from selflimited epilepsy with centrotemporal spikes (*n* = 3) or refractory focal epilepsy with neocortical spikes (*n* = 1) taken from (Feys et al., 2022). They were asked to sit still without moving while wearing 32 OPMs (Gen2, single-axis mode) on a flexible cap (number of channels kept for analysis: 29.25 ± 1.25). Recordings lasted 9.0 ± 1.9 min. The other patients were 2 adults (42 and 48 years) suffering from refractory focal epilepsy with medial temporal spikes taken from (Feys et al., 2025b). They were asked to sit still with eyes closed while wearing a flexible cap with 44 OPMs (27 Gen2 biaxial, 17 Gen3 triaxial; 40 kept for subsequent analysis, 19 Gen2 and 11 Gen3) for one patient, and 47 OPMs (27 Gen2 biaxial, 20 Gen3 triaxial; all kept for further analysis) for the other. Recordings lasted respectively 16 and 15 min. All IEDs in these data were originally marked by an experienced MEG clinician (number of detected events, 72.6 ± 78.4). We characterized these data using IED peak amplitude (maximum signal magnitude at IED peak time across all channels) and IED peak SNR (ratio of peak amplitude over baseline amplitude, measured as the SD of the same channel; baseline, 200 ms to 100 ms before peak) extracted from the IED response field (average of 600 ms epochs centered on peak time, excluding epochs intersecting bad time segments), as in the original works (Feys et al., 2022, 2025b). There, denoising prior to IED analysis relied on hard-threshold PCA with the number of components removed chosen after trial and error. This led to the removal of the first 3 components in the neocortical epilepsy dataset (Feys et al., 2022) and of the 15 first in the medial temporal epilepsy dataset where IEDs were noisier due to their deeper localization (Feys et al., 2025b). We compared results of this PCA-based IED analysis to results obtained using our calibrated SSB. Statistical comparisons of IED peak amplitude and SNR were performed with paired-sample *t* tests. The morphological similarity of the IED responses was assessed with canonical correlation (Feys et al., 2025b) and statistical effects with one-sample *t* test after Fisher transformation.

### 3.6 Investigating SSB in early neurodevelopmental data

Finally, we applied SSB to early neurodevelopmental OPM-MEG recordings where no sensor geometry measurement was available due to practical limitations of recording newborns and fetuses (Corvilain et al., 2025b, 2025a). We re-analyzed these challenging datasets using our pipeline based on general preprocessing steps and application of the calibrated SSB. Our goal was to compare these new results with previously published findings, which were obtained using several extra, arguably convoluted procedures using PCA (Corvilain et al., 2025b) or mixing PCA and regression analyses to mitigate the substantial noise afflicting these recordings (Corvilain et al., 2025a). Both datasets relied on an auditory paradigm similar to the vibration artefact experiment (section 3.4), i.e., sequences of 500 Hz pure tones, 100 ms long, including 15 ms rise and fall ramps to prevent auditory clicks.

#### Neonatal OPM-MEG recordings

(Corvilain et al., 2025b). The neonatal cohort was composed of 14 infants (7 females; mean 32.6 days ± 2.4 days). One participant was rejected because of technical difficulties. Subjects underwent the auditory paradigm (65 dB, 5 min with random interstimulus interval between 500 and 600 ms, 620 ± 108 trials) comfortably placed on their parent’s lap while wearing a small-size flexible cap equipped with 19 to 30 OPMs (27 ± 3, mean ± SD over subjects; 12 ± 8 Gen2 biaxial; 15 ± 9 Gen3 triaxial). ***Fetal OPM-MEG recordings*** (Corvilain et al., 2025a). The second cohort was composed of 21 pregnant women with uncomplicated pregnancies in their late third trimester of gestation (mean 32 years ± 3.3 years; gestational age, 38.3 weeks ± 1.1 weeks), of which two were rejected due to technical problems (abnormal noise floor). For fetal MEG recordings, participants were equipped with a stretchy maternity belt mounted with 45 to 54 OPMs (50 ± 3; 29 ± 2 Gen2 biaxial; 21 ± 2 Gen3 triaxial) and underwent the auditory paradigm (95 dB, 7 min, random inter-stimulus interval between 500 ms and 900 ms, 691 ± 104 trials). An accelerometer (3-axis 3G accelerometer, AALTO – Brain Research Unit) attached to the OPM belt was also used to record movements caused by the mother’s breathing synchronously to OPM-MEG.

In the original works (Corvilain et al., 2025a, 2025b), the general preprocessing steps described in section 3.2 were supplemented by several denoising steps. A more restrictive identification of bad time segments was performed automatically using a robust z-score rejection. In the neonatal OPM-MEG data, this was followed by a hard-threshold PCA rejection of the 3 largest principal components. In the fetal OPM-MEG data, the accelerometer signal was regressed out to reduce a large breathing artefact, along with the channels of a sub-selection of “external” sensors (i.e., all but the 32 most central OPMs on the belt most likely to record fetal brain activity) to suppress noise, followed by PCA rejection of the 3 largest principal components and a further regression of the vibration artefact at the level of the auditory ERF described below (for details, see (Corvilain et al., 2025a)). Our goal with these datasets was to compare the results obtained with this signal processing and those obtained with a straightforward denoising based on a calibrated SSB, bypassing these extra regression steps (and thus retaining all OPM channels, not just the central ones).

All subsequent analyses of auditory ERFs closely followed (Corvilain et al., 2025a, 2025b) for comparative purposes. Data were temporally segmented into epochs ranging from –100 ms to 700 ms around the start of sound presentation. Epochs intersecting bad time segments were excluded, interpolated linearly, and extended by 1.5 s after filtering, to avoid filtering effects. The numbers of trials used for ERF analysis were, for the neonatal data, 412.7 ± 104.8 with the original pipeline and 459.7 ± 110.5 with SSB; and for the fetal data, 465.5 ± 104.4 with the original pipeline and 520.1 ± 111.6 with SSB. The remaining epochs were baseline corrected (100 ms pre-stimulus time) and averaged within each participant. Finally, for both datasets, channels exhibiting a baseline temporal SD greater than 2 SDs above the average of all channels were marked as noisy and rejected. The neonate ERF dataset contained 27 ± 3 OPMs with the original pipeline (11 ± 7 Gen2 biaxial; 15 ± 9 Gen3 triaxial) and 27 ± 7 with SSB (11 ± 7 Gen2 biaxial; 15 ± 9 Gen3 triaxial); the fetal dataset contained 31 ± 1 OPMs with the original pipeline (12 ± 2 Gen2 biaxial; 19 ± 2 Gen3 triaxial) and 49 ± 4 with SSB (28 ± 2 Gen2 biaxial; 21 ± 2 Gen3 triaxial). Given the challenge of averaging ERFs across subjects with no consistent head positioning with respect to sensors and no sensor digitization, we summarized the auditory ERFs in terms of their dominant response curve (DRC) defined, for each subject, as the principal component of each individual ERF data restricted to the five most responsive channels. The scale of each DRC was readjusted to correspond to the root-mean-square of these five channels. We computed the grand average of these curves and their standard error of the mean (SEM). The response latency was calculated as the peak time of the DRC after a polynomial fit (to prevent picking up noise peaks; polynomial degrees chosen to minimize latency variation within each cohort, 15 for neonatal, 8 for fetal). To assess the impact of our SSB pipeline vs. the original pipeline, we statistically compared the DRC peak amplitudes and the DRC latency differences using paired-sample *t* tests.

## 4 Results

### 4.1 Calibration of SSB interference suppression

The effect of SSB on empty-room OPM-MEG recordings with a phantom head is depicted in Figure 1 as a function of increasing prospective values of the SSB threshold parameter *α* from zero up to 50%. Interference suppression with SSB reduced the average noise PSD in the 1–40 Hz frequency band whatever the parameter value, from 0.025 pT/√Hz (raw signal) to 0.017 pT/√Hz for *α* ≥ 1% (Figure 1A). The calibrated value was thus set to *α* = 1% according to our calibration heuristic. The clear denoising impact of SSB on OPM signal traces is illustrated in Figure 1B. Of note, the plateau in Figure 1A beyond the calibration value indicates that the gap hypothesis holds, i.e., larger regularization values do not further suppress activity in the complementary “signal subspace” (Theory section 2.5). The calibration heuristic at *α* = 1% was reproducible over time (Figure 1C) and in the fetal recording position (raw PSD, 0.024 pT/√Hz; SSB with *α* = 0%, 0.020 pT/√Hz; SSB with *α* ≥ 1%, 0.018 pT/√Hz).

**Figure 1:**
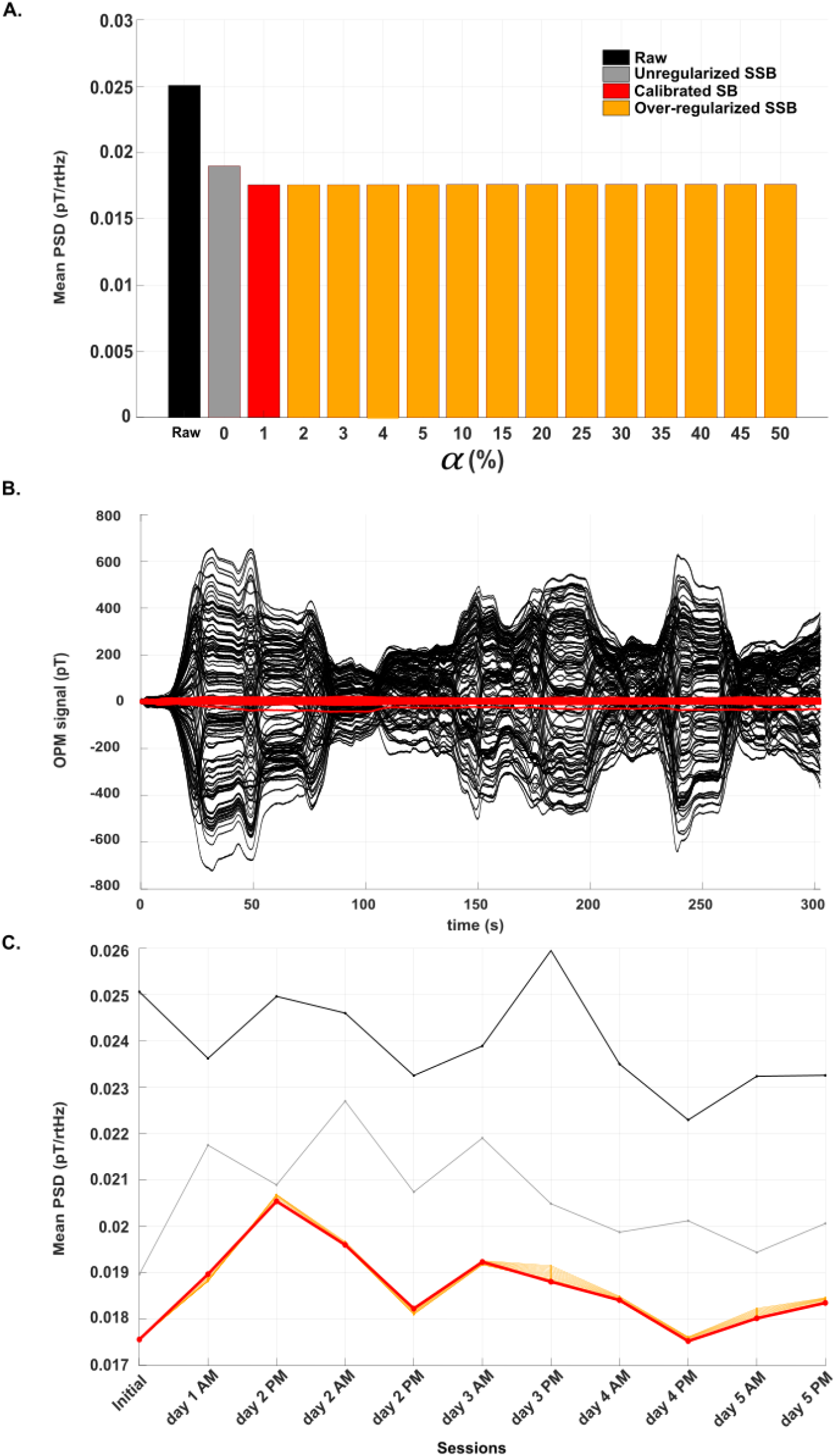
Calibration of SSB on empty-room OPM recordings. **A**. Bar plot of the 1–40 Hz PSD in the initial calibrating session for different values of the SSB threshold parameter α, measured as a percentage. Black, raw data without denoising; grey, SSB without regularization (α = 0%); red, SSB calibration heuristic (α = 1%); yellow, over-regularized SSB (α ≥ 2%). **B**. Corresponding time course of the raw data (black) and data corrected by the SSB with α calibrated at 1% (red). **C**. Time evolution of the 1–40 Hz PSD from the initial calibrating session (Figures 1A,B) to 5 days of empty-room recordings taking place 9 months later. The black curve corresponds to raw data and the others, to SSB at different values of the threshold parameter α (grey, 0%; red, 1%; yellow, ≥2%). The tight clustering of red and yellow curves confirms the stability of calibrated value α = 1% over time.

### 4.2 Denoising performance

We proceeded with further experiments probing the ability of SSB with its free parameter *α* calibrated at 1% to suppress the “interference subspace” covering typical OPM-MEG artefacts, and its ability to preserve the “signal subspace” covering neural activations. We started with the denoising of several simulated and physiological artefacts (Figure 2). The impact of SSB on OPM recordings in several noise conditions is summarized in Figure 2A.

**Figure 2:**
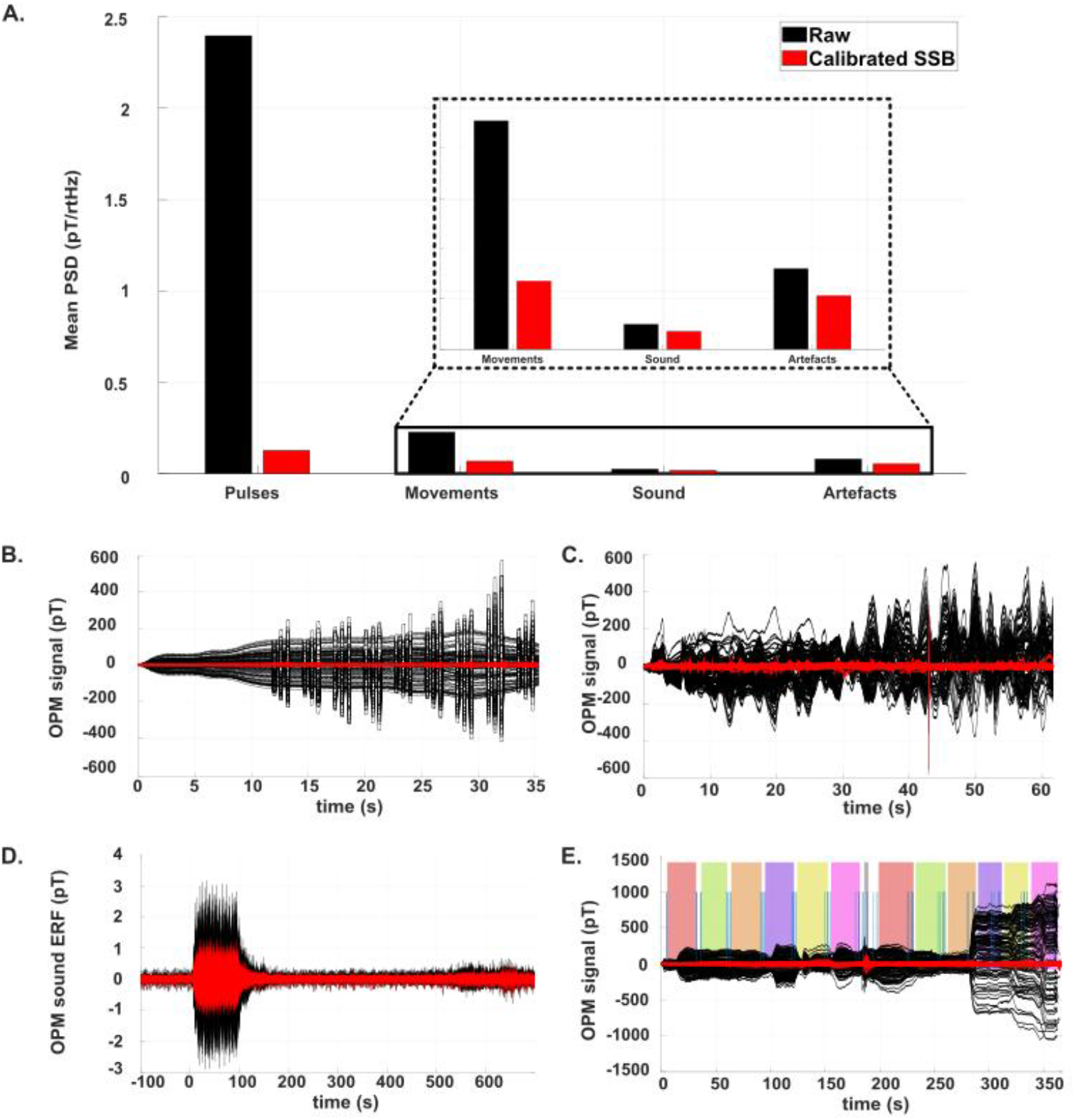
Denoising performance of calibrated SSB (red) compared to raw data (black). **A**. Bar plot of the mean 1–40 Hz PSD in different noise conditions: pulses (**B**), rigid movements (**C**), non-rigid sensor vibrations (**D**), and artefacts on a human subject (**E**). **B**. Time course of OPMs on phantom head while pulsing wall coils. **C**. Time course during rigid movements of the phantom head. **D**. Evoked response of OPMs vibrating due to external sound pulses. **E**. Time course of physiological artefacts of a human subject. Artefacts in order: blinking (light red box), lateral and then horizontal eye movements (light green box), contraction of jaw muscles (light orange box), clenching of teeth (light purple box), contraction of neck muscles (light pink box), and head movements (light gray box); sequence repeated twice (except for head movement).

#### Large field perturbations

Pulsing wall coils generated large magnetic fields (hundreds of pT, up to 580.5 pT) superimposed on the ambient magnetic drifts (Figure 2B, showing the pulsing of eight coils, each energized three times at three different intensities). Both the ambient and coil signals were strongly suppressed after SSB (largest pulse intensity reduced to 53.2 pT). Quantitatively, the mean 1–40 Hz PSD decreased more than tenfold, from 2.39 pT/√Hz to 0.13 pT/√Hz with SSB (Figure 2A). Of note, the noise floor after SSB still remained an order of magnitude larger than in the empty-room recording considered in Figure 1A (0.017 pT/√Hz), presumably because of the absence of active coil compensation in this experiment. ***Head movement artefacts***. Figure 2C shows the OPM recording on a phantom head under small rigid movements and the clear suppression of these low-frequency, large (up to 580.2 pT amplitude at one point) artefacts after SSB. Quantitatively, the noise floor decreased from 0.22 pT/√Hz down to 0.067 pT/√Hz after SSB (Figure 2A). The limited effect size on the noise floor, compared to the striking impact on the OPM traces in Figure 2C, is due to our restriction of the PSD to 1–40 Hz. ***Vibration artefacts***. The impact of sensor vibrations due to sound pulses led to comparatively small effects at the level of the continuous OPM-MEG signals, with a noise floor reduction from a modest 0.024 pT/√Hz without interference suppression down to baseline 0.017 pT/√Hz (Figure 2A). Figure 2D shows the ERF locked on the start of sound presentation, with a 3.0 pT oscillation during sound presentation (first 100 ms) clearly visible on the ERF traces and reduced to 1.3 pT after SSB. ***Physiological artefacts***. Finally, Figure 2E shows an OPM-MEG recording reproducing different types of human artefacts (blinking, lateral and horizontal eye movements, contraction of jaw muscles, clenching of teeth, contraction of neck muscles, and head movements). Here the impact of SSB on the 1–40 Hz noise floor was quite marginal (mean PSD reduced from 0.080 pT/√Hz to 0.053 pT/√Hz; see Figure 2A), contrasting with the clear effect seen in Figure 2E because the large-amplitude parts of these artefacts were very slow so that they were already strongly suppressed by simple frequency filtering above 1 Hz.

Taken together, these results show that SSB interference suppression successfully denoises OPM-MEG signals, with an added value compared to frequency filtering that depends on the dynamics of the artefact. As a purely spatial filter though, these experiments provide proof of concept that SSB does suppress a big part of the “interference subspace.”

### 4.3 Preservation of neural activity

It is also essential to assess whether the “signal subspace” is preserved. We used IEDs as model of clearly identifiable neural events, and compared their characteristics obtained after SSB to previous results obtained after PCA denoising (Figure 3).

**Figure 3:**
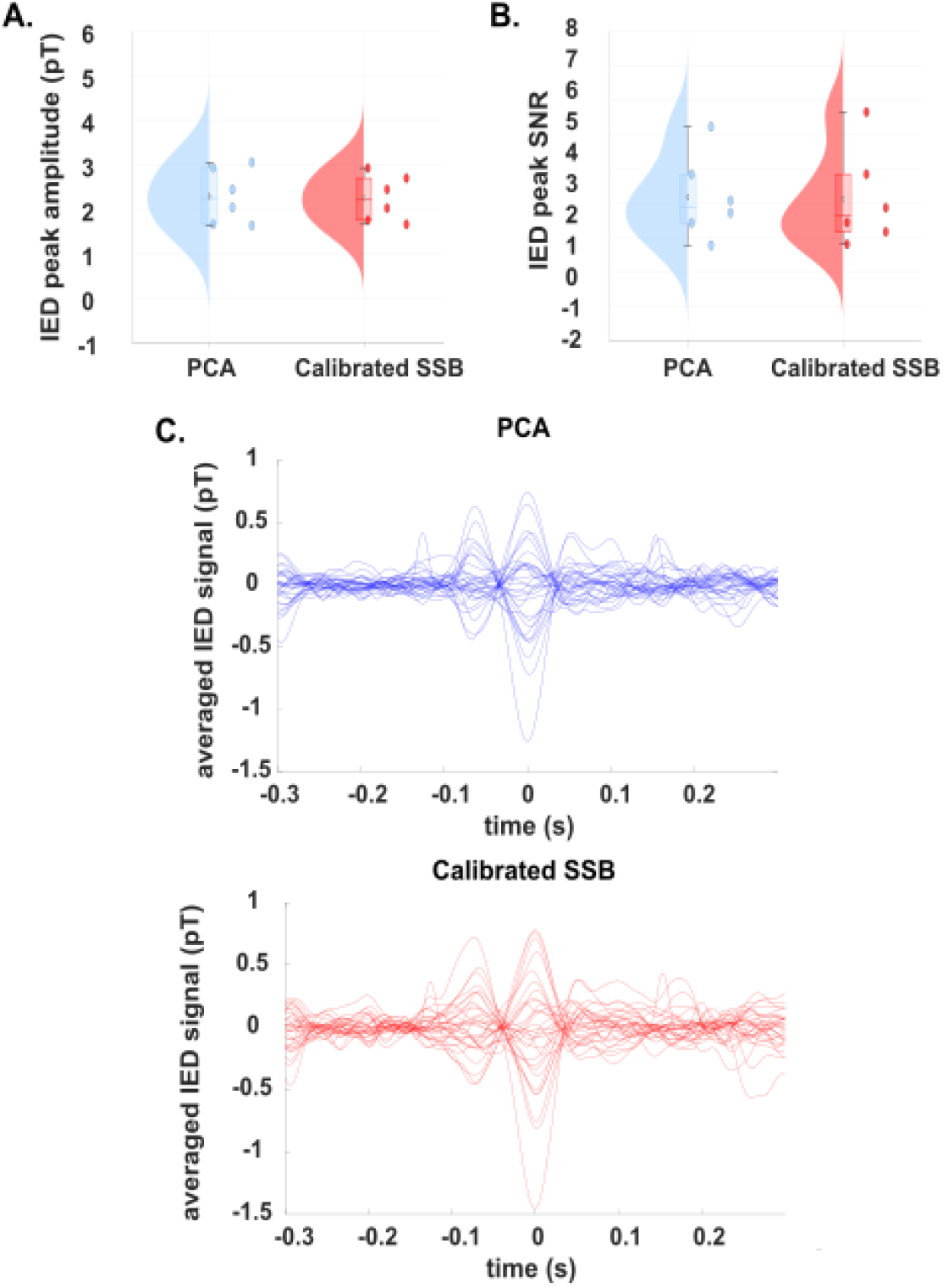
Preservation of IEDs by calibrated SSB (red), compared to PCA (blue) originally used (Feys et al., 2022, 2025b). **A**. Peak amplitude. **B**. SNR. **C**. Temporal morphology of the mean IED in a patient with neocortical epilepsy (for PCA, 3 principal components removed).

Figure 3A shows the distribution of the mean IED peak amplitude across the 6 patients following the originally published PCA denoising (Feys et al., 2022, 2025b) and interference suppression with the proposed calibrated SSB. The two approaches led to similar IED amplitudes (PCA, 2.0 pT ± 1.0 pT; SSB, 2.0 pT ± 1.0 pT; *t* test, *t*_5_ = 0.68, *p* = 0.52). Similarly, there was no difference in the distribution of IED peak SNR (Figure 3B; PCA, 3.18 ± 1.2; SSB, 3.13 ± 1.3; *t*_5_ = 0.35, *p* = 0.74). Figure 3C shows the IED temporal morphology in an example patient illustrating the similarity of both denoising techniques. This similarity was formally confirmed at the group level using canonical correlation testing (*R* = 0.75 ± 0.23; *t*_5_ = 3.78, *p* = 0.0064).

These results show that calibrated SSB performed equivalently to the original analysis by a clinical MEG expert; in particular, the morphology and amplitude of detected IEDs were similar. In other words, SSB did not suppress or deform IED events, suggesting that it preserves neural activity despite the cross-channel correlations.

### 4.4 Application to neurodevelopmental OPM-MEG

Finally, we tested our calibrated SSB with the most challenging datasets in which no information about sensor geometry was available—precisely the purview of SSB. We focused on the auditory responses in neonates as described in (Corvilain et al., 2025b) and fetuses as described in (Corvilain et al., 2025a). In both cases, we compared DRC characteristics (amplitude and latency) of auditory ERFs obtained after SSB to those previously published and obtained after dedicated preprocessing including PCA and several regression steps (Figure 4).

**Figure 4:**
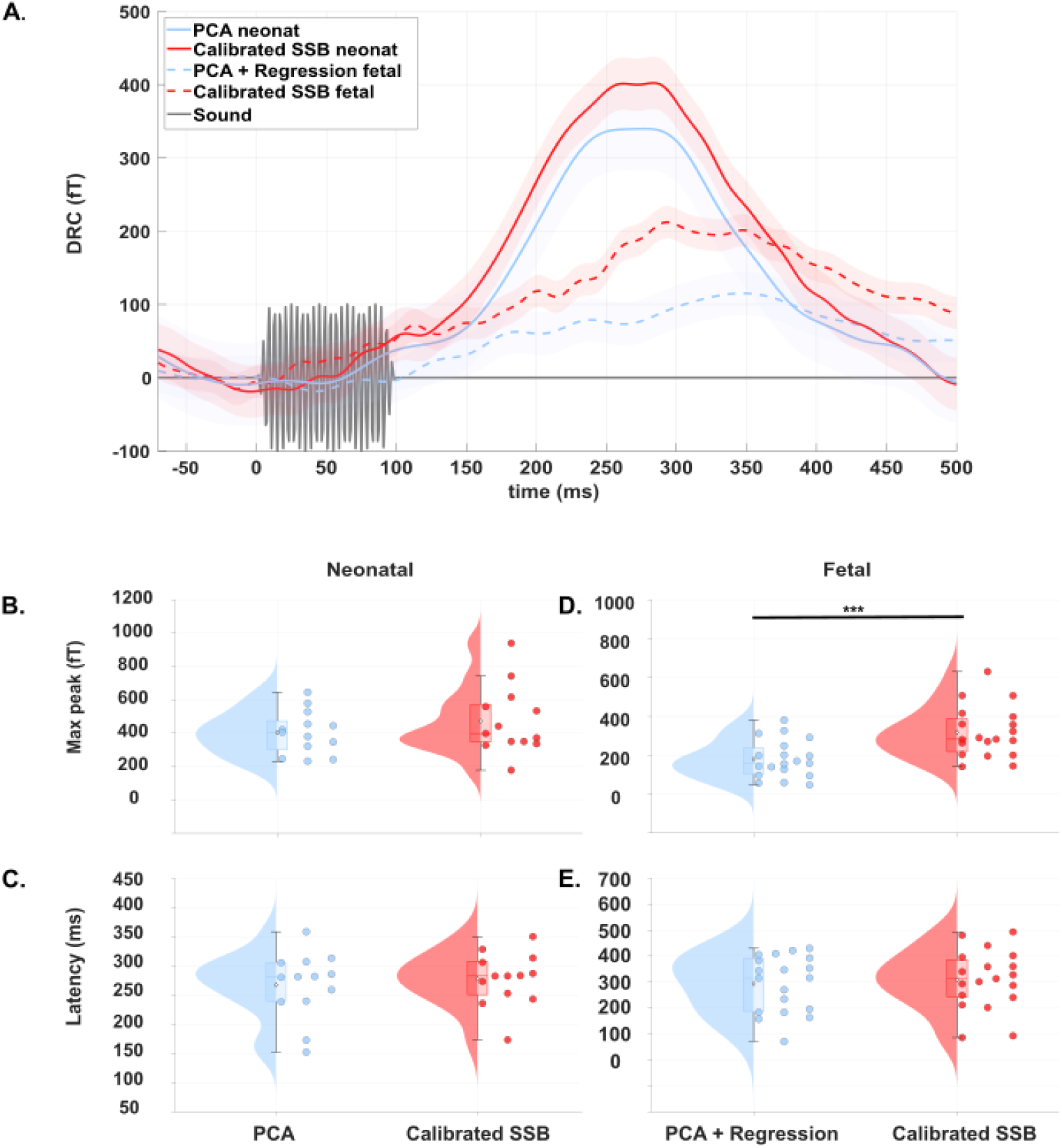
Auditory evoked responses in neurodevelopmental OPM-MEG obtained using calibrated SSB (red) and the original, PCA/regression-based pipeline (blue), for neonatal subjects (**A, B, C**) and fetuses (**A, D, E**). **A**. Group mean and SEM of auditory DRCs superimposed on the sound stimulus trace (grey, shown in arbitrary unit). Solid curves represent neonatal data; dotted curves represent fetal data. **B**,**D**. Comparison of peak amplitudes. **C**,**E**. Comparison of latencies. ***: p < 0.001.

#### Neonatal auditory responses

Figure 4A compares the DRC of the auditory ERF obtained after SSB to that reported in (Corvilain et al., 2025b). The DRCs appeared very similar in both cases. Quantitative analysis of peak amplitudes (Figure 4B) and latencies (Figure 4C) revealed no significant difference (SSB, 472.66 fT ± 203.34 fT at 277.94 ms ± 45.34 ms; original, 403.47 fT ± 129.40 fT at 267.41 ms ± 56.54 ms; *t* test, *t*_12_ = –1.96, *p* = 0.074 for amplitude, *t*_12_ = –2.15, *p* = 0.052 for latency). ***Fetal auditory responses***. In the case of fetal OPM-MEG data reported in (Corvilain et al., 2025a), the DRC obtained after SSB had a noticeably larger peak amplitude (Figure 4A). The peak amplitude was indeed significantly larger with SSB (318.39 fT ± 129.40 fT; Figure 4D) than with the original data (179.32 fT ± 95.93 fT; *t*_18_ = –5.33, *p* < 0.001) with no clear change in baseline activity, suggesting that SSB allowed to preserve more neural signal (Corvilain et al., 2025a). Latencies, on the other hand, were not significantly different (SSB, 309.15 ms ± 113.08 ms; original, 293.36 ms ± 108.52 ms; *t*_18_ = –2.02, *p* = 0.058; Figure 4E). Critically, the convergence of the fetal OPM-MEG responses processed with SSB and those processed with the original approach strengthens the evidence for a fetal neural origin of these responses.

## 5. Discussion

This paper introduces SSB—a software approach to OPM-MEG interference suppression based on a sensor-level, gain-preserving version of beamforming. We demonstrated how to calibrate SSB with a single empty-room OPM recording on a phantom head and showed that this procedure is reproducible over an extensive period of time. We provided proof of concept that the calibrated SSB suppresses the typical large-amplitude interferences that plague wearable MEG while preserving neural signals—in other words, that it works as a proper OPM-MEG interference suppression method. The key practical aspect that distinguishes SSB from landmark MEG signal denoising techniques is its freedom from prior measurements of OPM gain and geometry. This enables denoising OPM-MEG data in experimental situations where such measurements cannot be performed reliably, including prominent cutting-edge applications being actively pursued such as scalp MEG in difficult patients with epilepsy (Feys et al., 2023e; Feys & De Tiège, 2024), neonatal MEG (Chen et al., 2019; Rhodes et al., 2024), and fetal MEG (Dunn et al., 2015; Sheridan et al., 2010). We focused here on example datasets in each of these three domains and demonstrated that SSB leads to similar or better results compared with dedicated pipelines specifically developed for each situation, but without the need of fine-tuning. As such, SSB may provide a unifying framework for the preprocessing of OPM-MEG signals—again, at least when calibration of sensor gain and geometry is unreliable or unavailable.

### 5.1 Setting up SSB interference suppression in practice

The principles of SSB make it stand between SSS in the sense that it works at the sensor level and separates between a signal subspace and an interference subspace (Taulu et al., 2004), and source-space beamforming in the sense that denoising is based on the suppression of correlated activity (Hillebrand & Barnes, 2005; Van Veen et al., 1997). It is noteworthy that our SSB should not be confounded with the closely named “signal space separation beamformer” (Vrba et al., 2010), which differs fundamentally from the SSB in both aim and method as it implements a version of beamforming applied to SSS multipole coefficients. Conceptual aspects of SSB are discussed at length in the Theory section; we further consider here practical aspects of its implementation for OPM-MEG analysis. Setting up the SSB spatial filter requires two key choices: first, the input covariance matrix ***C*** dictating what “correlated activity” means; and second, the calibration of the free SSB parameter *α* dictating where is the separation between the “interference subspace” to be suppressed and the “signal subspace” to be preserved.

#### Choice of the input covariance

We focused here on the covariance matrix ***C*** derived from the raw OPM data. While this is the textbook choice in beamforming for MEG source projection (Sekihara & Nagarajan, 2008), other options have also been explored such as the common beamformer based on a covariance averaged across conditions (Lucena Gómez et al., 2021) or a non-adaptive beamformer based on resting-state covariance (Moiseev et al., 2015). None of these possibilities would work here though, as different recordings—even test/retest sessions of the same condition in the same subject—could be characterized by different artefacts, i.e., the SSB spatial filter weights must be constructed from the very data to denoise. Critically, what we mean here by “raw data” is that OPM signals are unfiltered, but importantly the exclusion of bad channels remains fully warranted for the proper estimation of input covariance (in complete analogy with the necessity of excluding bad channels before running SSS or performing MEG source reconstruction; see, e.g., (Wang et al., 2025a)). Alternatively, if one had priors on the frequency content of interferences, a possibility would be to extract the covariance after frequency filtering. Movement artefacts that dominate OPM-MEG signals are typically sluggish, so it would be reasonable to input the covariance of low-pass filtered data to the SSB spatial filter. This would *de facto* render the question (considered at length in this paper) of the preservation of neural activity by SSB moot, since high-frequency correlated patterns—including neural activity— would then be excluded from what the SSB considers as correlated activity to suppress. This is, in fact, one of the approaches that we considered while developing the method; however, denoising proved less efficient than with SSB built from the raw data covariance (data not shown). In hindsight, the reason is that many large low-frequency OPM-MEG interferences, such as movement artefacts, leak into higher frequencies and still dominate frequency ranges containing neural activity. In other words, there is no clear spectral gap separating the interference and the signal subspaces (hence the need for spatial filtering). Instead, the gap in amplitude proved much more reliable, leading us to the topic of SSB threshold calibration.

#### Calibration of the threshold paramete

Having established the choice of covariance, the next critical implementation step is the selection of the SSB threshold *α*, whose value effectively separates the space of correlated activities in terms of their relative amplitude. Here, we provided a heuristic calibration procedure that led to *α* = 1%, meaning that the SSB spatial filter weighted down modes of correlated activity that explained more that 1% of total input data variance, and that it preserved modes explaining less that 1% of variance. Importantly, while this heuristic worked successfully in our datasets and proved stable, we do not recommend taking this 1% threshold at face value when setting up SSB for the first time. This is because the *α* calibration might critically depend on the OPM-MEG system, the type of MSR, and its magnetic environment. Still, precise calibration might not be so critical; given the large amplitude gap that separates typical OPM-MEG interferences from their complement (see also Figure 1A), any reasonable choice for the SSB parameter *α* within that gap might work well enough.

Another critical remark is that SSB interference suppression does not fully liberate OPM-MEG from hardware denoising. The unit-gain constraint underlying SSB spatial filtering does not require the hardware needed for sensor gain calibration (such as recent “halo” calibrators, see (Hill et al., 2025)), but it still assumes that these gains are fixed throughout the experiment, i.e., without nonlinear gain changes induced by large magnetic fields (Tierney et al., 2019). This means that hardware reduction of the environmental field via passive compensation, MSR degaussing, and active compensation with coils (Boto et al., 2018; Holmes et al., 2019) all remain warranted for proper use of SSB interference suppression on OPM-MEG recordings. This being said, the development of OPM electronics operating in closed loop (Alem et al., 2023; Feys et al., 2023b; Schofield et al., 2024) might partially free OPM-MEG from the strict dependency on active compensation.

### 5.2 The purview of SSB—when to use it or not

Having discussed the question of how to set up SSB interference suppression in practice, we move onto the topic of when using SSB. The whole starting point for its development was to free OPM-MEG denoising from the calibration of sensor gain and geometry. We should make it clear though that, in our opinion, there is no doubt that the development of hardware solutions for the precise measurement of OPM channel gains, localization, and orientation—either based on external coils or bespoke calibrator systems (Hill et al., 2025; Iivanainen et al., 2022)—is ultimately the right direction to follow for the field of OPM-MEG at large. Fine calibration of sensor gain and geometry not only enables usage of the best, most principled interference suppression algorithms on the market such as SSS and extensions (Holmes et al., 2023a; McPherson et al., 2025; Taulu et al., 2004; Tierney et al., 2021, 2024), but it also renders MEG forward modeling and source localization techniques more precise (Hill et al., 2025; Steinsträter et al., 2010). Therefore, when technology allows, we recommend without ambiguity favoring sensor calibration-dependent methods. The purview of SSB is mostly restricted to “extreme” cases of OPM-MEG experiments where sensor gain and geometry cannot be measured or relied upon. This is somewhat analogous to the development of surrogate methods enabling OPM-MEG source localization in subjects or patients where anatomical magnetic resonance imaging cannot be obtained (Corvilain et al., 2025b, 2025a; Feys et al., 2023a; Rhodes et al., 2025).

Still, SSB interference suppression has a few advantages outside this realm of “extreme OPM-MEG.” Denoising based on SSS-like techniques does not only assume fine calibration of sensor gain and geometry but also sufficiently homogeneous and sufficiently dense spatial sampling (Holmes et al., 2023a), which to date remains a challenge with OPM-MEG. Technically, SSS relies on OPM-MEG systems working in the oversampling regime (Seymour et al., 2022), which requires not merely increasing OPM channel count using multi-axis magnetometers but also the sheer number of OPMs (Wens, 2023). The recent development of high-density OPM-MEG systems with 64 (Schofield et al., 2024) or even 128 OPMs (Alem et al., 2023) helps closing in to the oversampling regime (Wens, 2023). Notwithstanding these technological advances, SSB is comparatively less sensitive to sensor coverage and density (to a limit at least) and, crucially, can be used with non-conventional OPM montages with partial (Feys et al., 2022) or inhomogeneous (Feys et al., 2023d) coverage of the scalp, as occurs, e.g., in cases of simultaneous OPM-MEG/SEEG recordings (Feys et al., 2025a). Likewise, OPM coverage in peripherical nerve (Bu et al., 2022), spinal cord (Mardell et al., 2024), and gut recordings (Liang et al., 2024) strongly depart from the sphericity assumed by SSS, in which cases SSB may provide an efficient alternative to denoising. Finally, geometry-dependent techniques also assume that the relative position of sensors is rigidly fixed throughout the recording, which neglects possible non-rigid sensor movements when using flexible caps for clinical OPM-MEG (Feys et al. 2022) or stretchable belts for fetal OPM-MEG (Corvilain et al., 2025a). We showed here that SSB can deal with realistic nonrigid deformations, albeit only with partial success, as demonstrated with the incomplete suppression of the sensor vibration artefact. It is possible that this interference was not sufficiently correlated across sensors to enable full suppression by SSB. On the other hand, in fetal OPM-MEG data, non-rigid deformations of the belt due to respiration were successfully removed by SSB, which allowed bypassing the accelerometer regression step.

It is also noteworthy that a wide array of alternatives to geometry-dependent SSS-like denoising algorithms exists besides SSB. The most popular examples comprise the regression of reference signals (Adachi et al., 2001), PCA (Barbati et al., 2004), and ICA (Vigario et al., 2000); for systematic reviews, see (Seymour et al., 2022) and (Wang et al. 2025). Here, we explicitly compared SSB to previous OPM-MEG preprocessing relying on PCA (to which SSB relates as a soft-threshold, gain-preserving version; see Theory section 2.4) and regression pipelines. We found that SSB performed similarly but without the need of specific fine-tuning of denoising parameters. The main added value of SSB is that a single, all-purpose calibration step effectively replaces both the user-dependent selection of how many principal components must be removed (in our OPM-MEG epilepsy data) and the experiment-dependent selection of what reference signals must be regressed out (in our fetal OPM-MEG data). The scope of ICA is typically quite different and restricted to physiological MEG artefacts such as eye blinks and heartbeats, rather than the large artefacts that dominate OPM-MEG signals. In fact, ICA does not bide well with these large interferences because they do not follow the assumption of positive excess kurtosis underlying blind signal separation by ICA (Hyvärinen & Oja, 2000) whereas physiological noises do (Vigario et al., 2000). That is why ICA is generally used after an initial pass at MEG denoising (notwithstanding some instances where ICA was used as initial denoising, though these applications were limited to cryogenic, fixed-array MEG systems where interferences are orders of magnitude smaller than in moving OPM-MEG; see (Mantini et al., 2008)). Conversely, SSB successfully suppresses—by design—dominating OPM-MEG interferences thanks to their well-defined gap in amplitude (see above), but not physiological artefacts because they cannot be clearly isolated from neural signals based on their amplitude. From the viewpoint of SSB, physiological noise presumably belongs to the “signal subspace.” Therefore, SSB is no substitute for ICA or other refined MEG denoising tools; rather, its scope is simply to render OPM-MEG signals tractable for these tools.

In conclusion, the purview of SSB interference suppression is to denoise OPM-MEG recordings in extreme experimental situations where advanced technological solutions for MEG calibration cannot be used reliably. In this paper, we had in mind two distinct examples of “extreme OPM-MEG” that we illustrated here. First, there is the domain of OPM-MEG epilepsy, one of the leading clinical applications of cryogenic MEG, where patient compliance may not allow for accurate calibration of OPM gain and geometry (Feys et al., 2023e). Second, there is the domain of early neurodevelopmental OPM-MEG, an innovative direction of OPM research plagued by practical recording difficulties (Corvilain et al., 2025b, 2025a). In our view, the application to fetal OPM-MEG data provides the best illustration of the usefulness of SSB interference suppression. Not only did we showcase the ability of SSB to dig up subtle fetal brain responses buried in complex artefacts (non-rigid deformations due to mother breathing, sensor vibrations, etc.) with no calibration of OPM gain and geometry available, but our analysis also provided confirmation of the neural origin of these responses first described in (Corvilain et al., 2025a). Looking further ahead, the enhanced flexibility of the OPM technology makes it easy to envision the emergence of innovative applications of OPM-MEG in the future. It will be interesting to see how SSB— alongside hardware technological developments—will help getting noisy OPM-MEG recordings closer to the theoretical limits of their potential (Wens, 2023).

## Acknowledgements

M.F. and P.C. have been supported by the Fonds Erasme (Brussels, Belgium; research convention “Alzheimer”). M.F. is supported by a research grant from INNOVIRIS (Brussels, Belgium; research convention “DetectDem”) and C.C. by a grant from the Fonds de la Recherche Scientifique (F.R.S.–FNRS, Brussels, Belgium; research convention T.0026.24, attributed to J.B.). O.F. and L.F. acknowledge financial support from the Fonds pour la Formation à la Recherche dans l’Industrie et l’Agriculture (FRIA, F.R.S.–FNRS). X.D.T. is clinical researcher at F.R.S.–FNRS. Data acquisitions were supported by the F.R.S.–FNRS (research project T.0026.24, J.B.; research credit J.0043.20F, X.D.T.; equipment credit U.N013.21F, X.D.T.; Audacious Medical Grant Neuro 2024, X.D.T.) and the Fondation Jaumotte-Demoulin (Brussels, Belgium; attributed to J.B.). The OPM-MEG project at the HUB–Hôpital Erasme is financially supported by the Fonds Erasme (research convention “Les Voies du Savoir” and Clinical Research Project, X.D.T.).

## Author contributions

V.W. conceptualized study. M.F., P.C., O.F., C.C., L.F. contributed to data acquisition. M.F., P.C., V.W. developed methodology. M.F., P.C. wrote codes. M.F., P.C., O.F., C.C., J.B., X.D.T, V.W. analyzed data. M.F., V.W. wrote initial manuscript draft. J.B., X.D.T. provided resources and funding. All authors reviewed and approved final version of submitted manuscript.

## Ethics

Epilepsy OPM-MEG studies were approved by the HUB–Hôpital Erasme Ethics Committee (P2019/426/B406201941248 and P2020/664). Patients and their legal representatives (for minors) provided written informed content prior to inclusion. The HUB–Hôpital Erasme Ethics Committee approved the experimental protocol of the neurodevelopmental OPM-MEG experiments (P2021/141/B406202100081). Parents/mothers gave informed consent prior to testing and received monetary compensation for their participation. All analyses were carried out in accordance with approved guidelines and regulations.

## Data and code availability

Clinical epilepsy data will be made available upon reasonable request to X.D.T. after approval of institutional authorities (Université libre de Bruxelles & Hôpital Universitaire de Bruxelles). The neurodevelopmental dataset is publicly available at https://doi.org/10.5281/zenodo.10213856.

Code of the SSB interference suppression, alongside analysis scripts for the different datasets, will be made publicly available upon publication.

